# Efficiency of a Randomized Confirmatory Basket Trial Design Constrained to Control the False Positive Rate by Indication

**DOI:** 10.1101/2020.07.22.216127

**Authors:** Linchen He, Yuru Ren, Han Chen, Daphne Guinn, Deepak Parashar, Cong Chen, Shuai Sammy Yuan, Valeriy Korostyshevskiy, Robert A. Beckman

## Abstract

**PURPOSE:** Molecular oncology determines biomarker-defined niche indications. Basket trials pool histologic indications sharing molecular pathophysiology, potentially improving development efficiency. Currently basket trials have been confirmatory only for exceptional therapies. Our previous randomized basket design may be generally suitable in the resource-intensive confirmatory phase, maintains high power, and provides nearly *k*-fold increased efficiency for *k* indications, but controls false positives for the pooled result only. Since false positive control by indications (FWER) may sometimes be required, we now simulate a variant of this basket design controlling FWER at 0.025*k*, the total FWER of *k* separate randomized trials.

**METHODS:** The previous design eliminated indications at an interim analysis, conducting a final pooled analysis of remaining indications. To control FWER, we rechecked individual indications at a prospectively defined level of statistical significance after any positive pooled result. We simulated this modified design under numerous scenarios varying design parameters. Only designs controlling FWER and minimizing estimation bias were allowable.

**RESULTS:** Sequential analyses (interim, pooled, and post-individual tests)) result in cumulative power losses. Optimal performance results when *k* = 3,4. We report efficiency (expected # true positives/expected sample size) relative to *k* parallel studies, at 90% power (“uncorrected”) or at the power achieved in the basket trial (“corrected”, because conventional designs could also increase efficiency by sacrificing power). Efficiency and power (percentage active indications identified) improve with higher percentage of initial indications active. Up to 92% uncorrected and 38% corrected efficiency improvement is possible, with power ≈ 60%.

**CONCLUSIONS:** Even under FWER control, randomized confirmatory basket trials substantially improve development efficiency. Initial indication selection is critical. The design is particularly attractive when enrollment challenges preclude full powering of individual indications.

## INTRODUCTION

Molecular oncology has led to increasingly numerous biomarker-defined niche indications.^1^ For example, lung cancer now includes several small subgroups with distinct therapies. This may lead to clinical trial enrollment challenges, and to additional development expense and delay. Conversely, a targeted therapy may have potential application in numerous indications, as well as in multiple combination settings. We previously discussed the need for increased development efficiency in design of proof of concept studies and associated Go-No Go decisions, given the large number of potential hypotheses worthy of testing.^2^ These issues also apply to the resource-intensive confirmatory phase of development, contributing to the very high cost of therapy, as well as to decisions not to develop drugs for niche indications. Exclusive reliance on traditional development, one indication at a time, may be unsustainable.

Master protocols^3,4^ can potentially increase the efficiency of drug development, and facilitate development of niche indications. Master protocols include platform trials, in which different therapies are perpetually cycled through an ongoing trial, resulting in notable operational efficiencies; umbrella trials, in which multiple therapies are matched to multiple biomarker subgroups within a single traditional organ-system-based indication, and basket trials.

In a basket trial, traditional indications are grouped together in a basket based on a shared molecular or pathophysiologic characteristic thought to predict utility of a therapy. These indications may borrow information from each other, or may be frankly pooled, leading to large improvements in development efficiency. The challenge in optimizing basket trials comes from the risk of heterogeneity between indications despite shared biomarkers sometimes conferring similar benefits. Highly active indications may carry inactive indications along with them to create a positive pooled result. Conversely, inactive indications may dilute the effectiveness of active indications, leading to a negative pooled result.

Several efficient basket trial approaches use response rate data in the exploratory setting. Most of these are based on Bayesian hierarchical models^5–7^ although one considers the likelihood that the indications all come from one statistical distribution, versus the likelihood they are best modeled individually.^8^

The first oncology basket trial in a regulatory approval setting was for imatinib, which had already demonstrated extraordinary value in chronic myelogenous leukemia, and had been rationally designed based on considerable scientific evidence.^9^ The study was non-randomized and based on response rate, with very small sample sizes. Forty indications were evaluated in less than 200 patients. One approval resulted from 1 response in 5 patients. Related designs resulted in approvals for transformational drugs designed for alterations in b-raf and ntrk oncogenes.^10,11^ Recently, the immune checkpoint inhibitor pembrolizumab was approved in multiple solid tumors, based on a basket trial pooling patients from these indications with a DNA repair defect resulting in microsatellite instability.^12^ In all of these cases, the drugs and biomarkers were supported by unusually strong scientific evidence and achieved transformational results in underserved indications, and thus were able to merit approval based on single-arm response rate data in relatively small populations.

### Motivation for the Research

In contrast, many effective oncology drugs have required rigorous randomized designs in the confirmatory setting. We developed a confirmatory basket trial design^13,14^ which, in addition to being applicable in a single-arm fashion or with response rate endpoints, can also be utilized in a randomized setting using time to event (TTE) endpoints such as progression free survival (PFS) and overall survival (OS). Randomization is generally important in approval of agents whose clinical benefit must be measured based on TTE endpoints, unless the effect size is transformative. This design was intended to have potential suitability for the majority of confirmatory settings.

The original adaptive design resembles a funnel (Figure 1A, Methods). Indications are carefully selected, and then filtered (removed) in several “pruning steps”, first with data external to the study (i.e. maturing phase 2 data from the same drug, data from other agents in the class), and then with data from an interim analysis, which may be based on a surrogate endpoint considered predictive of the definitive endpoint (i.e. PFS predicting OS) or on early analysis of the definitive endpoint. After pruning, a sample size adjustment may be applied to the remaining indications, which are then pooled. The study concludes with a pooled analysis of the remaining indications based on the definitive endpoint, and is positive if statistically significant benefit is shown.

**Figure 1.**
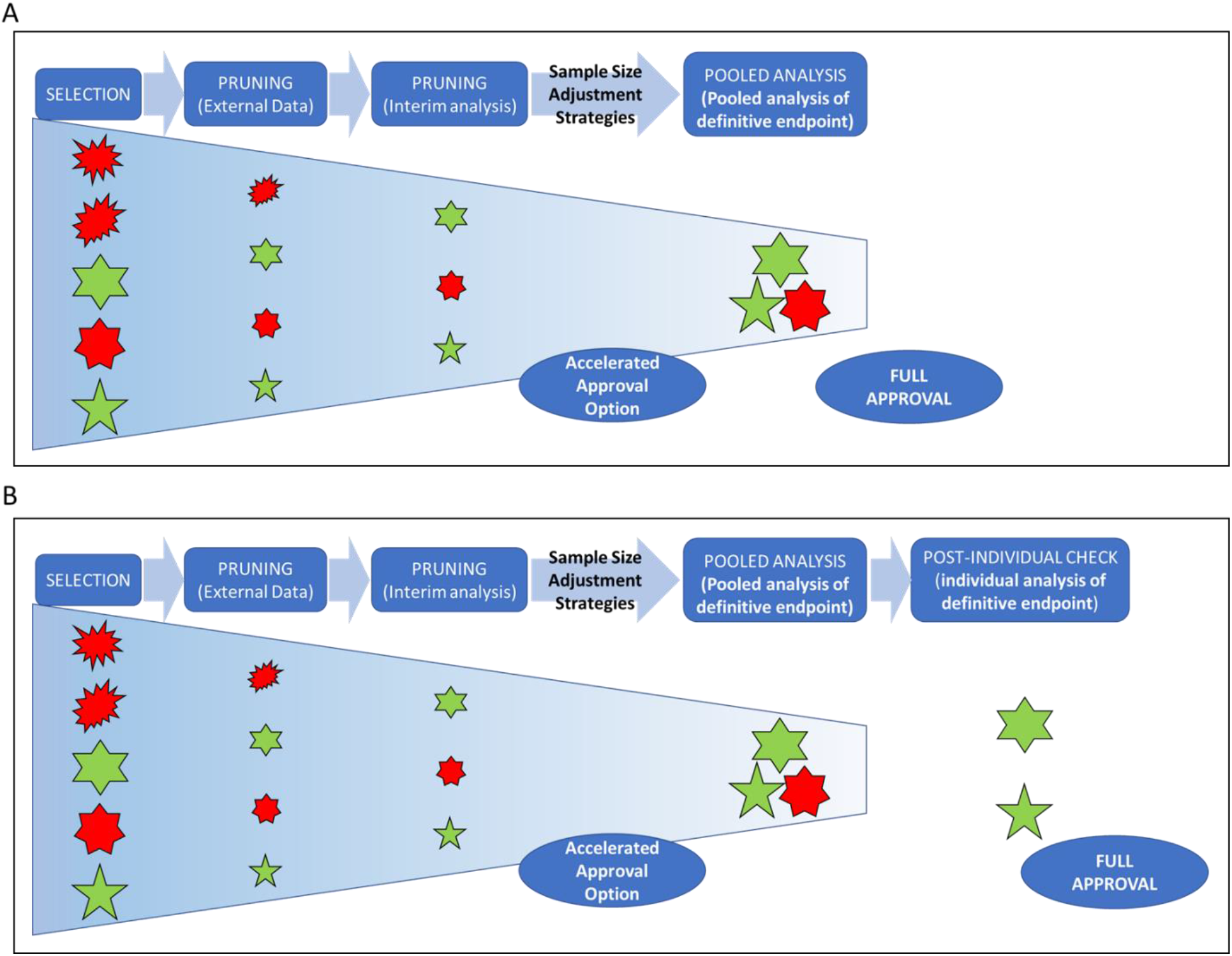
Confirmatory basket trial design (A) with pruning and pooling as in previous studies^13,14^, and (B) with pruning, pooling, and post-individual check simulated in this study. The designs (1) conduct a basket trial that consists of *k* tumor indications; (2) prune (remove) indications based on external data; (3) conduct an interim analysis independently for each tumor indication. Indications that meet these criteria may in some cases be eligible for accelerated approval, indications that do not are pruned; (4) adjust sample size of remaining indications as needed; (5) conduct a pooled analysis of the remaining indications; (6) in the current design (B) only, conducts a prospectively defined post-individual check for each indication involved in the pooled analysis. In previous studies (A), indications that passed the pooled analysis may potentially be eligible for full approval, whereas in the new design utilized in this study (B), indications must also pass a simultaneous post-individual check to be potentially eligible for full approval.

This design demonstrated a dramatic increase in development efficiency, which we define in this work as (expected number of true positives/expected sample size), and maintained acceptable power over a variety of scenarios even with inactive indications in the basket.^13,14^ However, it was designed to control the false positive rate only in the pool as a whole, but not by traditional indication subgroups (family-wise error rate, FWER) (Figure 2A, Methods, Supplemental Methods). In this paper, we define active and inactive indications as indications in which the test drug does or does not provide clinical benefit, in the unknown state of nature. FWER by indication subgroup, simply called FWER in this paper, is defined as the probability that one or more inactive indications will be approved by the design. Control of the FWER may be recommended by health authorities in confirmatory settings.^15^

**Figure 2.**
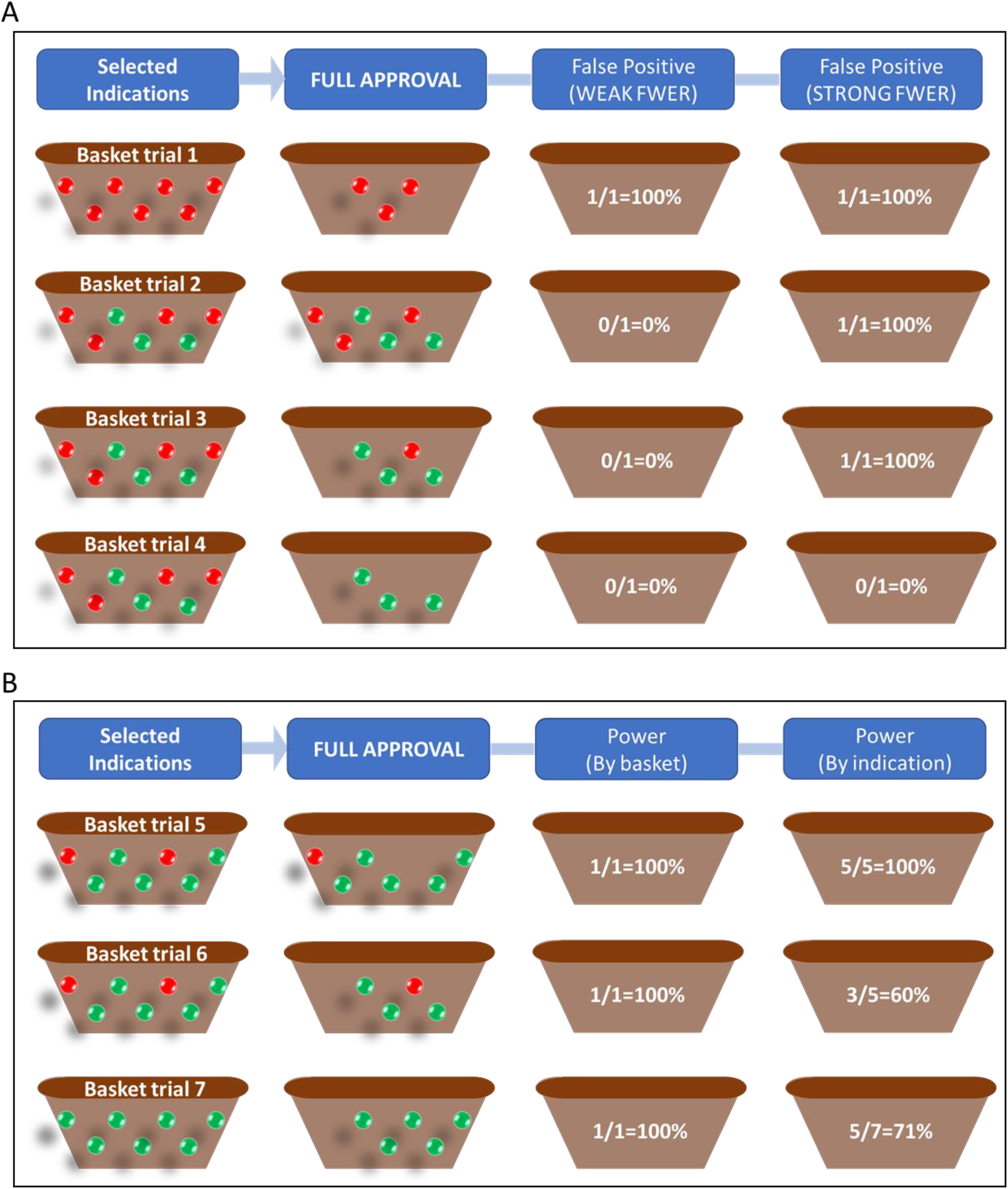
Measurements of false positive rate (type I error) (A) and power (B) showing the difference between criteria used in this study and less stringent criteria. Examples are shown for illustration. Each row represents an example and marbles represent the active (green) and inactive (red) indications. The left-hand column represents the initial selected indications. The second column represents the approved indications. The third column represents traditional criteria, and the fourth column represents the more stringent criteria used in this study. (A) This study is designed to control the family-wise error rate (FWER) by indication subgroup rather than false positive rate in the pool in the case where all indications are inactive. In the third column, a false positive is scored only when a basket is approved containing only inactive indications. In the fourth column, a false positive is declared if a single inactive indication is approved. (B) Power results are evaluated by indication in this study rather than by basket. Considering the power by basket (third column), the approved basket can be considered as a true positive if it has at least one active indication. In the fourth column, the power by indication is defined as the percentage of active indications approved, and can be calculated as 100%, 60%, and 71% for the three examples, respectively.

Cunanan et al documented that a well-known exploratory basket trial design did not control FWER, and, without opining whether this was acceptable in the exploratory setting, advocated for disclosure of performance properties for complex designs.^16^ We agree, and characterized our original randomized confirmatory basket trial in this respect, finding that it did not control FWER with acceptable power levels (data not shown). It may still be suitable in those confirmatory settings where control of FWER is not recommended.

In order to control FWER, and also maintain acceptable power levels, we implemented a modification of the initial design in which, whenever the pooled result is positive, each indication is re-checked at low to medium stringency for statistical significance before final approval (Figure 1B, Methods). The series of tests, each at lower stringency (higher alpha) is sufficient to control FWER to a prespecified level more stringent than each of the individual checks. As has been shown in biological systems where high fidelity with minimum energy expenditure is required^17,18^, repeated testing at lower stringency is more efficient than a single highly stringent test.

We describe the performance of this modified design in extensive simulations, focusing on a scenario with the same TTE endpoint at interim and final analyses. By varying design parameters (Table 1, Methods, Supplemental Methods), we created numerous design variants. Acceptable design parameters must control FWER to the same level as a system of individual randomized studies for each indication in parallel, i.e. approximately 0.025 multiplied by the number of indications. Further, acceptable design parameters must not introduce bias of greater than 10% in the effect size estimate, and the estimated 95% confidence interval of the effect size must cover at least 90% of simulation runs. For the input parameters to be considered acceptable, these constraints must be met regardless of the number of inactive indications within the basket.

**Table 1.** Glossary of Terms

| Design parameters | Descriptions |
| --- | --- |
| The state of nature |  |
| $g$ | The number of active indications at the beginning |
| $\theta_i$ | The hazard ratio of experimental arm vs. control arm for the definitive endpoint for indication $i$ |
| Study design input parameters |  |
| $k$ | The number of tumor indications at the beginning |
| $D$ | Sample size adjustment strategies |
| $\alpha_t$ | A common bar for the interim analysis to prune inactive indications |
| $\alpha$ | False positive rate for the pooled analysis |
| $\beta$ | False negative rate for the pooled analysis |
| $\alpha_{post}$ | A common bar for the post-individual tests |
| Outcome measurements |  |
| $\alpha_{net}$ | The probability of the basket trial passing the pooled test and at least one false positive indication passing the post-individual test for a given value of $g$ |
| Family-wise error rate (FWER) | The maximum probability of the basket trial passing the pooled test and at least one false positive indication passing the post-individual test for any $g$. The possible values of $g$ indicate the “family” for which FWER is defined. |
| Power by indication | The probability of an active indication passing the interim test, the pooled test, and the post individual test |
| Power by basket | The probability of a true positive basket (with one or more active indications) passing the pooled analysis |
| Efficiency | The ratio of average number of active indications that pass the post-individual tests divided by the average sample size |
| Uncorrected relative efficiency | The efficiency of the basket design divided by that of a group of parallel traditional confirmatory studies, powered at 90%, investigating the same indications. |
| Corrected relative efficiency | The efficiency of the basket design divided by that of a group of parallel conventional studies at the same power, investigating the same indications. |

For design parameters meeting these constraints we characterize development efficiency and power as estimated by simulation. We define development efficiency as the expected number of true positive indications identified divided by the expected sample size. “Uncorrected” relative development efficiency is the efficiency of the basket design divided by that of a group of parallel traditional confirmatory studies, powered at 90%, investigating the same indications. As we do not achieve 90% power by indication in the basket trial, we present a “corrected” relative development efficiency adjusting for the power losses. As enhanced development efficiency can be achieved in conventional designs by reducing power alone^2^, corrected relative development efficiency compares the efficiency of the basket design to parallel conventional studies at the same power.

Power is evaluated by indication (Figure 2B), a more stringent criterion than traditional confirmatory studies, where subgroups are usually not formally powered. Power is therefore the fraction of active indications expected to be qualified for approval by the basket trial. For comparison, we also present the traditional power of the pooled analysis.

We present in Results selected designs that optimize corrected relative efficiency, either subject to a minimum power constraint (recommended development scenarios) or irrespective of power. These designs are characterized as a function of the number of total indications, the number of active indications, and the degree of activity (hazard ratio 0.5-0.8).

We discuss overall utility of the design and key learnings for performance optimization. We propose criteria for when control of FWER should and should not be recommended by health authorities in confirmatory studies. Finally, we outline future research topics aimed at addressing other potential concerns with randomized confirmatory basket trial designs.

## Methods

### Study Design Overview and Design Parameters (Table 1, Supplemental Table S1)

Consider a randomized confirmatory basket trial of an experimental therapy that consists of *k* tumor indications. For each indication, we perform 1:1 randomization (experimental vs. control), with a TTE variable as the primary endpoint of interest and *n* events per indication.

Figure 1B presents the current study design. We assume an interim analysis is conducted at a common information time *t* ∈ (0,1) based on *nt* events for all tumor indications, which we assume is also the same actual time. The study designer should choose sample sizes to make this approximately true, but may need to conduct several interim analyses and sample size adjustments in practice. We assume a common bar *α_t_* (one-sided nominal Type I error rate) across tumor indications for simplicity. A tumor indication is “pruned” from the study if it does not meet the bar for pooling. Remaining indications proceed to the pooled analysis. After the interim analysis, we adjust sample size for remaining tumor indications to account for lost sample size due to pruning. We consider three sample size adjustment designs^13,14^:

1. Design one (D1): No sample size adjustment.
2. Design two (D2): Aggressive sample size adjustment to replace all originally planned events in the pruned indications.
3. Design three (D3): Moderate sample size adjustment to replace future originally planned events after the interim analysis in the pruned indications.

Although endpoints for pruning and pooling may be different, in this work, herein we consider the same endpoints only. Denote (*α, β*) as the false positive and negative rates for one-sided hypothesis testing in the pooled population. The adjusted false positive rate *α*\* to control the false positive rate for the pooled analysis at the desired level will be calculated for each strategy with respect to a global null hypothesis that all indications are inactive (Figure 2A, Supplemental Methods)^13,14^. To control FWER, when a basket passes the pooled analysis, a prospectively defined individual post-pool analysis, examining each of the *m* tumor indications remaining in the pool after all pruning is complete, is conducted, termed a “post-individual test”. We also assume a common prospective post-individual bar *α_post_*, varied independently of *α_t_*. Indications that survive the post-individual test may be eligible for full approval. The three successive filters, i.e. pruning, pooled testing, and post-individual testing give a net false positive rate of *α_net_* corresponding to a given set of input parameters. FWER is considered controlled to the level of *α_target_* if *α_net_* ≤ *α_target_* for a given set of input parameters and all *g* from 0 to *k*.

### Outcome Measurements (Table 1, Supplemental Table S1)

There is no clear analog of conventional false positive and negative rates for basket trials. We denote the number of active indications by *g* ≤ *k*. We simultaneously consider the complete “family” of all scenarios involving *g* = 0,1,…, *k*. The familywise error rate (FWER) is defined as the maximum probability of at least one false positive indication passing a post-individual test over all possible values of *g*. The power is defined as the proportion of active indications passing the post-individual tests.

We also examine the efficiency, defined as the ratio of average number of true positive indications passing the post-individual tests divided by the average sample size. Other measurements include the coverage of the confidence interval (CI) of the hazard ratio (HR) and bias of the estimated HR. In evaluating outcome measurements, we threshold the FWER at the level 0.025*k*, the coverage of the 95% CI of the HR as greater than 90% of the simulation runs, and the bias of estimated HR less than 10%. Design parameter combinations which cannot meet these criteria irrespective of the value of *g* in our simulation are not allowed. Finally, we examine power by indication and efficiency for each allowed design and utilize a ratio to compare efficiency relative to a reference design.

The reference design for the uncorrected relative efficiency calculation assumes parallel, independent Phase III designs planned for each indication with the false positive and false negative rates (*α_ref_, β_ref_*) set to (0.025,0.1).

For calculation of relative efficiency corrected for power losses, we first determine the power by indication of a basket study with the same design parameters and inputs, then set *β_ref_* equal to 1 minus this power, while maintaining *α_ref_* at 0.025, and finally proceed as for the calculation of relative efficiency above.

In our simulation, we assume that for each indication the true hazard ratio (HR) *θ_i_*; *i* = 1,…, *k* is either at a null value *θ*_0_ = 1 or at an effective value *θ_a_* = 0.5 – 0.8, which is constant across indications within a scenario. We vary *k* from 3 to 6. We fix *α* = 0.025 and explore different values of *β, α_t_* and *α_post_*. For each setting, we generate 10000 simulated trials. Simulations and analysis are performed using R (version 3.6). Source code and details of simulation settings and outcome measurements are available in Supplemental R codes.

## RESULTS

We explore power and efficiency with various input parameters (Table 1, Supplemental Table S1). We control FWER at FWER ≤ 0.025*k* for all values of *g*, the number of active indications, from 0 to *k* inclusive. We provide recommended optimal design parameters and associated performance results (Figure 3, Supplemental Figure S1). These recommendations consider both corrected relative efficiency and power. We also provide design parameters that optimize corrected relative efficiency irrespective of power (Figure 4, Supplemental Figure S2) and show associated performance results. All simulation results are available in Supplemental Table S2.

**Figure 3.**
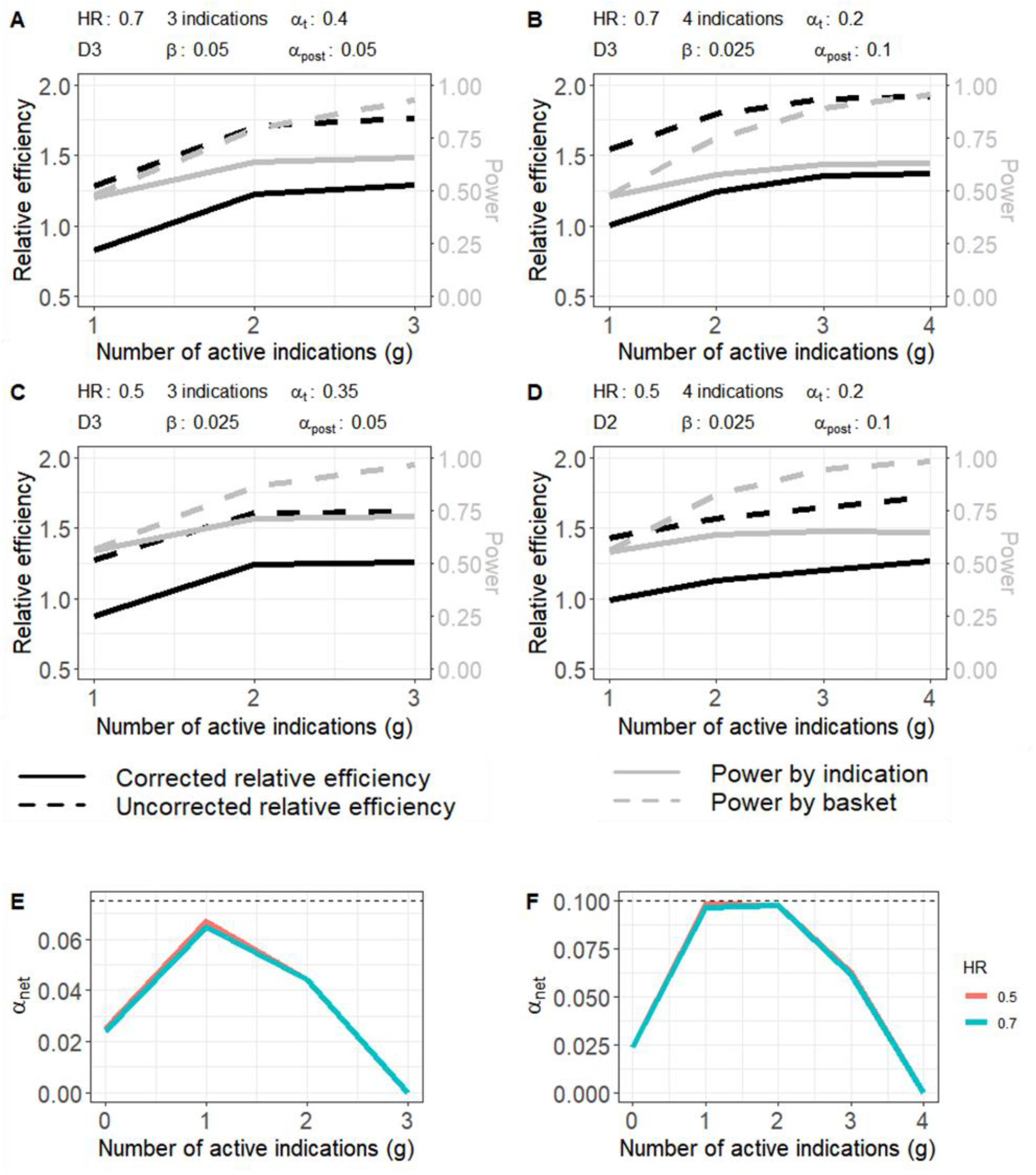
Recommended development approaches for (A) 3 indications with *HR* = 0.7, (B) 4 indications with *HR* = 0.7, (C) 3 indications with *HR* = 0.5, and (D) 4 indications with *HR* = 0.5. The X-axis represents the number of active indications (indications in which the drug provides clinical benefit), the primary Y-axis (left) represents the uncorrected/corrected relative efficiency, and the second Y-axis (right) represents the power by indication and by basket. (E) *α_net_* is shown on the y axis, and the number of active indications on the x axis, for *HR* = 0.5 (orange) and 0.7 (blue). *α_net_* is controlled at ≤ 0.025fc (dotted horizontal line) for k =3. (F) same as (E), for *k* = 4.

**Figure 4.**
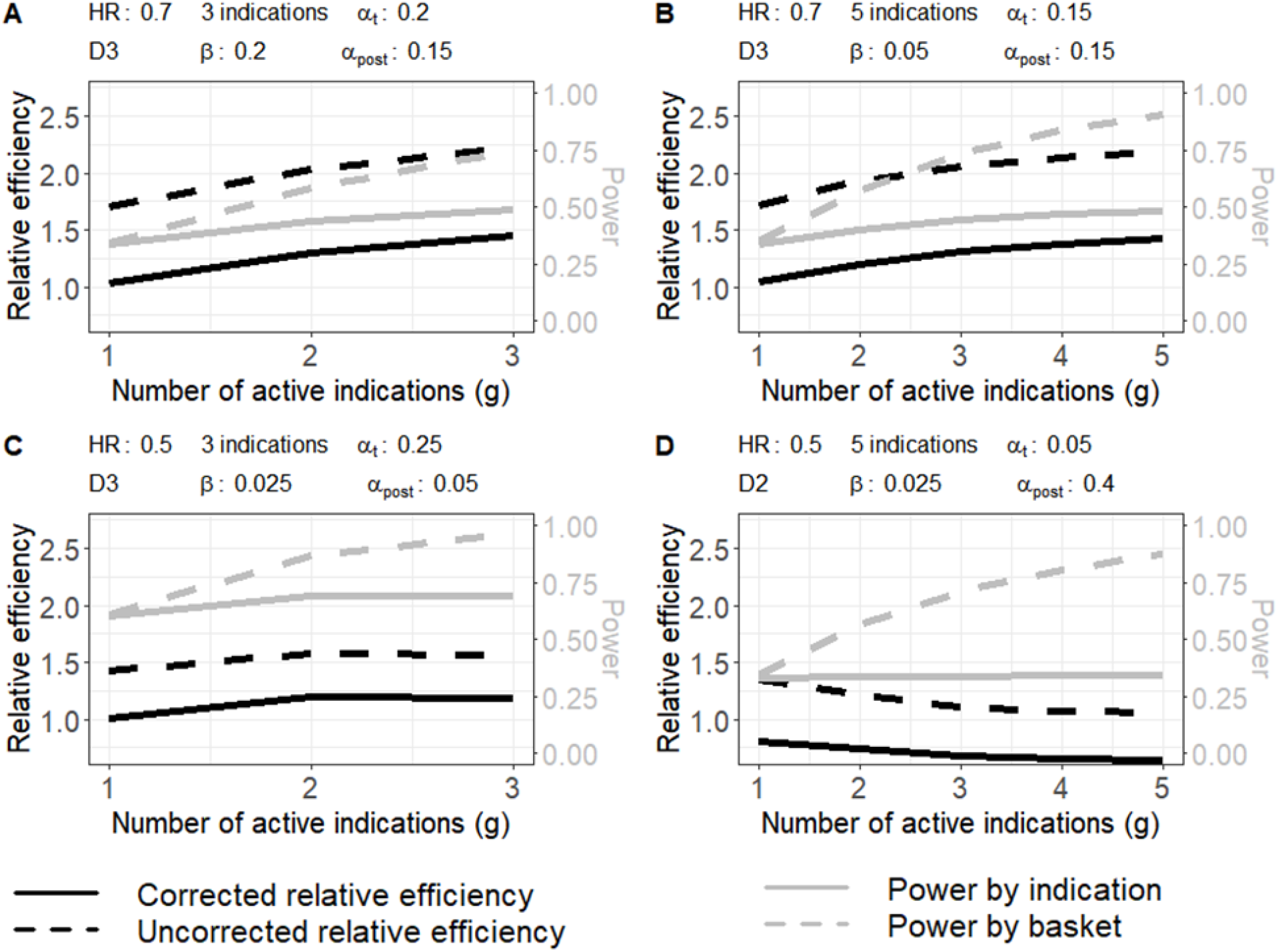
Cases with maximum corrected relative efficiency for (A) 3 indications with *HR* = 0.7, (B) 5 indications with *HR* = 0.7, (C) 3 indications with *HR* = 0.5, and (D) 5 indications with *HR* = 0.5. The X-axis represents the number of active indications (indications in which the drug provides clinical benefit), the primary Y-axis (left) represents the uncorrected/corrected relative efficiency, and the second Y-axis (right) represents the power by indication and by basket.

### Decreased power and/or efficiency if the majority of indications are inactive

Power and/or efficiency increases with the proportion of indications that are active. (Figures 3–4, Supplemental Figures S1-S2). Performance deteriorates when multiple inactive indications are present in the basket.

As the hazard ratio of positive indications increases, indicating a decreased therapeutic effect size, the corrected relative efficiency of the recommended development scenario generally increases, but the power of the recommended development scenario generally decreases. This pattern suggests a tradeoff between power and efficiency.

Maximal corrected relative efficiency is seen at *k* = 3,4.

In addition, FWER is more difficult to control with increasing number of tumor indications and increased therapeutic effect size, reflecting greater heterogeneity (Supplemental Table S2). False positive rate *α_net_* is generally maximal when *g* is approximately half of *k*, again reflecting maximal heterogeneity.

### Recommended development scenarios

We have previously emphasized that this design requires careful initial indication selection.^13,14^ Users should strive to include only zero or one inactive indications in the study (Discussion). To determine recommended development scenarios, we considered these upside scenarios, specifically requiring power by indication greater than 60% when all indications are active. After implementing this filtering criterion in addition to the requirements for control of FWER, estimation bias, and confidence interval coverage at all values of *g*, only the scenarios with *k* = 3,4 remain. Figure 3 and S1 summarize the scenarios with optimal upside corrected relative efficiency for *k* = 3,4 and different values of the hazard ratio. Power by basket increases from 50% to greater than 90% for any scenario as the number of active indications increases from 1 to *k*, while the power by indications increases to 63% – 72% when all indications are active, and 62% – 71% when there is one inactive indication. For the uncorrected relative efficiency, scenarios with 4 indications exhibit about 43% – 92% efficiency improvement depending on g, 23% – 78% efficiency improvement for three indications. Corresponding ranges of corrected relative efficiency improvement are −17% – 29%, and −1% – 38%, for *k* = 3,4, respectively. Note if most of the indications are inactive, the design is inefficient. When all indications are active, or only one indication is inactive, the uncorrected efficiency improvement ranges from 61% – 78% and 65% – 92% for 3 and 4 indications, respectively, while the corrected improvement in efficiency ranges from 21% – 29% and 20% – 38% for 3 and 4 indications, respectively.

### Optimal corrected relative efficiency scenarios

Without filtering for power, Figures 4 and S2 summarize scenarios with maximal corrected relative efficiency for *k* = 3 – 5 and different values of the hazard ratio. Scenarios with *k* = 6 have inferior performance (Supplemental Table S2) and are not recommended.

Comparing to the recommended development scenarios, the optimal corrected relative efficiency scenarios present higher efficiency and lower power. Power by basket ranges from 34% – 98%, 26% – 98%, and 21% – 98%, depending on *g*, for *k* = 3,4,5, respectively. Corresponding power ranges for power by indication are 34% – 71%, 25% – 65%, and 20% – 59%. Uncorrected relative efficiency improvement ranges from 25% – 125%, 47% – 163%, and 6% – 172% depending on the value of *g* for *k* = 3,4,5 respectively. Corresponding ranges of corrected relative efficiency improvement are −9% – 47%, −6% – 67%, and −36% – 66%. When all indications are active or only one indication is inactive, the uncorrected efficiency improvement ranges from 46% – 125%, 65% – 163%, and 6% – 172%, for *k* = 3,4,5, respectively, while the corresponding corrected relative efficiency improvement ranges from 14% – 47%, 20% – 66%, and −36% – 66%. Removing the constraint on power increases relative efficiency gains, again indicative of the tradeoff between power and efficiency. In most cases, the basket trial design improves the efficiency, but this is not always the case (Figure 4D).

## DISCUSSION

Confirmatory basket trials potentially provide remarkable improvements in drug development efficiency. With pooling, a fold-improvement in efficiency comparable to the number of indications is possible while retaining high power for the pool, and controlling alpha with respect to a global null hypothesis.^13,14^

We studied a modification of the randomized confirmatory basket design^13,14^ in a more rigorous setting where control of FWER by indication subgroup is recommended. We have further studied the most challenging scenario, i.e. a slowly maturing TTE endpoint without a highly predictive surrogate. Performance characteristics for innovative designs should be publicly disclosed in detail, including in challenging settings, but this is not always the case.^16^ Under the conditions of the simulation, up to 92% improvement in relative efficiency, or 38% improvement corrected for reduced power by indication, is still possible, while controlling the false positive rate at a rate comparable to an equal number of parallel single-indication studies. Power of the pooled analysis (“power by basket”) remains high, and estimation bias and confidence interval coverage may also be well controlled. Power by indication can be as high as 60% – 73%, but declines if there are two or more inactive indications in the basket. The results apply only to the conditions of the simulation. As there are an infinite number of cases to simulate, a sponsor would need to reach agreement with health authorities on the range of scenarios to be simulated, as has been articulated previously for complex innovative designs.^19^

A study of ten oncology drugs found an average cost of $648 million for their clinical development, much of which spent in the resource-intensive confirmatory phase.^20^ Expense and prolonged clinical development time delays availability of therapies, and contributes to their high cost and potentially to unequal treatment access, major public health issues. We believe the confirmatory basket design or modifications thereof could contribute to the solution of these problems. Moreover, in oncology niche indications, this design could make drug development economically feasible even in the absence of a transformational therapy. Finally, transformational approaches such as immunotherapy must be further optimized, creating a broad universe of potential combination studies across populations sharing common characteristics such as high tumor mutation burdens. These therapy combinations may lead to important but incremental improvements that require a randomized TTE approach for confirmation.

We have previously suggested that the performance of randomized confirmatory basket trials depends on careful indication selection.^14^ This is even more true when control of FWER is recommended, as is documented in the results. These designs are inappropriate for a collection of miscellaneous uncharacterized indications. Any proposed indication should be supported by preclinical studies, and Phase 2 clinical and biomarker data.^14^ It may be helpful to filter these indications in Phase 2 with one of several exploratory basket trial designs^5–8^, especially if the indications have enrollment challenges. Real world data/evidence, especially concerning off-label use, may be of value in screening potential indications.^21,22^

The optimal range for indication number is narrow. For recommended drug development scenarios, which seek optimal corrected relative efficiency with a minimum constraint on power by indication, either 3 or 4 indications are optimal. If only corrected relative efficiency is optimized, one may also consider 5 indications. Too few indications and one does not get the benefit of a basket trial. Too many, and compensating for the large number of potential heterogeneity scenarios involved in controlling FWER becomes challenging. Interestingly, earlier work determining an optimal indication number in exploratory basket trials also recommended 3-5, despite a very different scenario and approach to optimization.^23^

Optimal stringency of filtering varies, but generally greater stringency is applied in post-individual tests than at interim. Higher stringency is perhaps better applied to more mature datasets. Optimal sample size adjustment (SSA) strategies varied. Sometimes moderate SSA (D3) was better than aggressive SSA (D2). Aggressive SSA may optimize power at the expense of overall development efficiency.^24^ The optimal nominal power for the pooled analysis also varies according to the power/efficiency tradeoff. Higher power requires disproportionate investment, as previously shown for proof of concept studies.^2^

It is important to consider if and when strong control of FWER by indication subgroup should be recommended for a confirmatory basket trial with pooling, an area of debate which will likely evolve. A conventional phase 3 study is really quite heterogeneous with respect to both known and unknown subgroups (Figure 5), and we do not control FWER in assessing the vast majority of these subgroups, employing other approaches.^25, 26^ In a basket trial, we invert the usual classification: molecular subgroups are now indication-defining, and organ sites, formerly indication-defining, are now subgroups. This alone may present a perceptual barrier to full acceptance of the concept. Collignon et al^15^ argue that homogeneity of outcomes may be difficult to interpret clinically since the populations are “different”, illustrating the unproven perception that differences in organ sites are more fundamental than the many other known and unknown differences between subpopulations. But they also provide a definition of subgroup homogeneity in striking agreement with our thinking: “homogeneous if they share important clinical characteristics such that, in light of the available scientific evidence, the interpretation of treatment effect and the assessment of benefit/risk are meaningful for the overarching target population…” This comes down in the end to science and medicine, not statistics, and indeed the scientific justification for pooling must be robust if we are to forego FWER control by subgroup. We know that even in the classic case of an antagonist of a driver mutation in the b-raf gene, drug effectiveness still depends on organ site.^10^ Increasing understanding of how driver gene mutations interact with tissue-specific gene expression programs may be important. If we do not have a robust justification for pooling, control of FWER by indication will be necessary. Collignon et al^15(SI)^ evaluate our original confirmatory basket design^13,14^, and our assertion that evidence supporting a consistent benefit/risk assessment across traditional indications should be provided at a level prospectively agreed with health authorities.^14^ They state that availability of post-approval data would be important. This should become more practical as electronic health records systems improve.

**Figure 5.**
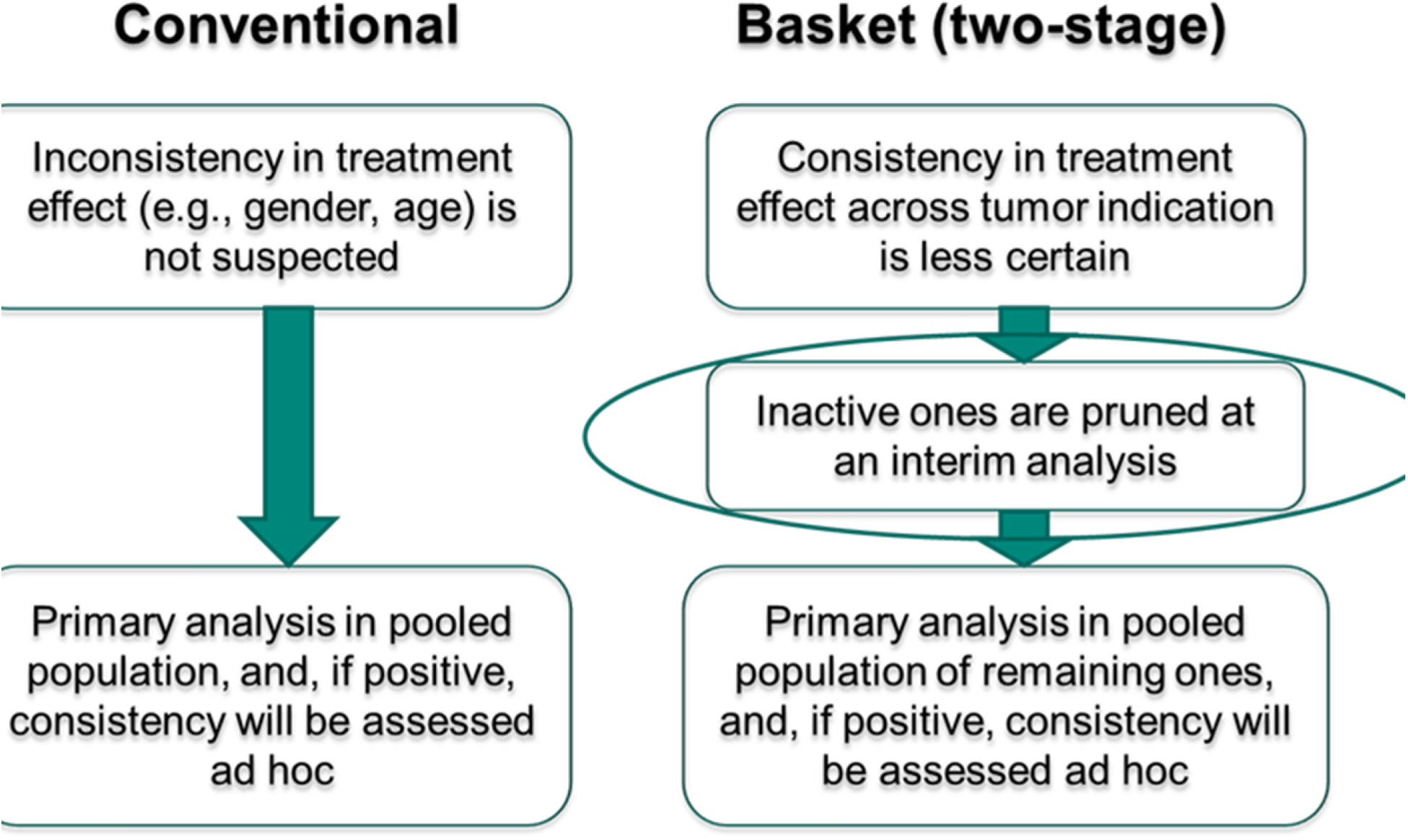
Parallels between a conventional phase 3 study (left) and the randomized confirmatory basket trial design considered in previous work without strong control of the false positive rate by subgroup (right). Both designs formally test a hypothesis for a main group and do not formally test hypotheses involving subgroups. In the conventional design, organ site is the defining characteristic of the main group, and both known and unknown subgroups are present, the former perhaps undergoing more informal subgroup analyses. In the randomized confirmatory basket trial, the traditional organ site classification is only a known subgroup, and a biomarker or other pathophysiologic feature defines the main group. Organ site classification is subjected to informal analysis only.

In ongoing research, we are considering important questions regarding the suitability of these designs for the confirmatory phase, in particular how to deal with differences between indications in endpoints and in safety issues. In future work, we will consider the effects of different control therapies, enrollment rates, and endpoint maturation rates. We are investigating real world data/evidence in indication screening and parameter estimation in simulations. Finally, current performance may be improved further by application of techniques previously devised for exploratory trials.^5–8, 27^

## ACKNOWLEDGEMENTS

We thank Drs. Zoran Antonijevic, Lisa LaVange, Martin Posch, and William Rosenberger for careful and insightful review of the manuscript.

