## Supplemental Materials for "Efficiency of a Randomized Confirmatory Basket Trial Design Constrained to Control the False Positive Rate by Indication"

### SUPPLEMENTAL METHODS

#### Overview of Type I error evaluation

In this basket trial setting (Figure 1B) designed to examine multiple indications ($k$ in number), there is no clear analog of conventional Type I error rates, since the possibility of committing a type I error may occur for tests of each indication. We consider the familywise error rate (FWER) by indication, which is defined as the probability of at least one false positive indication getting approved irrespective of the number active and inactive indications, defined respectively as indications in which the drug provides or does not provide clinical benefit in the unknown state of nature. This FWER considers a family of null hypotheses in which one or more of the $k$ indications are inactive, i.e. $2^k - 1$ null hypotheses.  The FWER is progressively controlled by three successive pruning steps, none of which individually provide type I error control as stringently as all three in sequence. Considering a basket trial that consists of $k$ tumor indications, these three analyses must be passed for an indication to be approved:

1. Interim analysis: For each of $k$ tumor indications, an interim analysis prunes (removes) inactive indications. We assume a common bar $\alpha_t$ (in terms of the one-sided nominal Type I error rate) for all tumor indications for simplicity.
2. Pooled analysis: For all remaining indications that pass the interim analysis, a pooled analysis is performed relative to the null hypothesis that all indications in the pool are inactive. The adjusted nominal level $\alpha^*$ is used (further details in next section)[13,14].

3. Post-individual check: For each indication that passes the pooled analysis, a prospectively defined post-individual check determines whether an indication may be eligible for full approval. We assume a common bar $\alpha_{post}$, which is varied independently of $\alpha_t$.

The three analyses are conducted at the nominal levels $\alpha_t$, $\alpha$*, and $\alpha_{post}$, however, none of these nominal levels quantifies the Type I error of the entire trial. Rather than control the false positive rate of any of three tests at the level of $0.025$ per indication, we evaluate the FWER of the entire trial according to the cumulative effect of three sequential tests. To evaluate the FWER, we design comprehensive simulation studies, where a range of nominal levels $\alpha_t$ and $\alpha_{post}$ is selected (Supplemental Table S1) and the adjusted nominal level $\alpha$* is calculated for each strategy. For a given set of design parameters and hazard ratio of active indications, the overall type I error resulting from the cumulative effect of the 3 steps for a given value of $g$ is $\alpha_{net}(g)$. We then vary $g$ from 0 to $k$ with all other parameter constant to get the corresponding value of FWER:

$$FWER = max\{ \alpha_{net}(g): g = 0, 1, \dots k\}$$

Based on simulations, we find design parameters under which the FWER is controlled at $\leq \alpha_{target} = 0.025k$ (see RESULTS).

##### The adjusted nominal level $\alpha$* in the pooled analysis[13,14]

The pooled analysis aims to examine that there is treatment effect in at least one tumor indication. This analysis combined with the previous pruning and sample size readjustment steps is controlled at type I error of $0.025$ for the global null hypothesis that all indications are inactive using methods from reference 13. Note that the definition of

Type I error of the entire trial in reference 13 is different from the FWER in this study. Specifically, in reference 13, let $Y_{i1}$ be the standardized test statistics based on the endpoint used for pruning at the interim analysis, and $Y_{i2}$ be the standardized test statistics based on the endpoint for pooling for the $i$-th tumor indication at the final analysis ($i = 1, \ldots, k$). Suppose that $m$ tumor indications are included in the pooled analysis ($m \geq 1$). Let $V_m$ be the corresponding standardized test statistics pooled from $Y_{i2}$, which can be written as $(\sum_{i=1}^{m} Y_{i2})/\sqrt{m}$. Denote α as the probability of incorrectly declaring activity in a basket when there is no treatment effect in any of the tumor indications. Chen et al.[13] formulated the probability of $V_m$ being statistically significant at the adjusted $\alpha^*$ level given $m$ out of k indications are in the pool, as

$$Q_0(\alpha^*, \alpha_t, m) = c(k,m)\, P_{H_0}\big(\cap \{Y_{i1} > Z_{1-\alpha_t}; i = 1, \ldots, m\}, V_m Z_{1-\alpha^*}\big)(1-\alpha_t)^{(k-m)}$$

where $c(k,m) = k\,!/\big((k-m)!\,m!\big)$ is the number of choices for selection of $m$ tumor indications from $k$. The overall Type I error probability with respect to the global null hypothesis after the pooled analysis, set at $\alpha = 0.025$, is $\sum_{i=1}^{k} c(k,m) Q_0(\alpha^*, \alpha_t, m)$. The adjusted level $\alpha^*$ , at which the pooled analysis is nominally set, can be solved based on the correlation between $Y_{i1}$ and $V_m$, $corr(Y_{i1}, V_m)$, considering the following three sample size adjustment strategies[13]:

1. Design one (D1): Sample size for each tumor indication is fixed upfront at *n* as planned. After pruning, the sample size will be less than or equal to the originally planned $kn.$ Under D1, $corr(Y_{i1}, V_m) = \sqrt{t/m}$. This strategy corresponds to no sample size adjustment.

2. Design two (D2): Sample size for each tumor indication will increase after the interim analysis so that the total sample size in the pooled analysis remains $kn$, which is greater than the sample size under D1 after pruning. Under D2, $corr(Y_{i1}, V_m) = \sqrt{t/k}$. This is the most aggressive sample size adjustment strategy.

3. Design three (D3): Sample size for each tumor indication will increase after the interim analysis so that the total sample size in the overall study remains $kn$. Thus, the total sample size in the pooled analysis is greater than the sample size under D1 but smaller than the sample size under D2. Under D3, $corr(Y_{i1}, V_m) = \sqrt{t/\left(mt + k(1-t)\right)}$.

**Simulation study**

In our simulation study, we used parameter values that are summarized in supplemental Table S1. We assume that for each indication the true hazard ratio (HR) $\theta_i; i = 1, \ldots, k$ is either at a null value $\theta_0 = 1$ or at an active value $\theta_a = 0.5$, 0.6, 0.7, or 0.8 and the true number of active indications is $g = 0,1, \ldots, k$. We fix $\alpha = 0.025$ and vary design parameters $k$ $(3,4,5,6)$ and $\beta$ $(0.025,0.05,0.1,0.2)$. Consequently, the total sample size in the pooled population ($kn$) can be calculated as $kn = 4\left(Z_{1-\alpha} + Z_{1-\beta}\right)^2/[(log\theta)^2 k]$. For simplicity, we set a common information time for the interim analysis $t = 0.5$, so that each indication should accrue $nt$ patients for interim analysis. We note that the in practice the indications should be chosen so that they are projected to reach $nt$ events at approximately the same time to avoid the operational inconvenience of multiple interim analyses and sample size adjustments. Furthermore, the sample size for each indication at pooled analysis should accrue as follows:

1. $\mathrm{n}_{i2} = \mathrm{n}(i = 1, \dots, m)$ under D1;
2. $\mathrm{n}_{i2} = \frac{k\mathrm{n}}{m}(i = 1, \dots, m)$ under D2, which is greater than the sample size under D1;
3. $\mathrm{n}_{i2} = \mathrm{n}\left(t + \frac{k(1-t)}{m}\right)(i = 1, \dots, m)$ under D3, which is greater than the sample size under D1 but smaller than the sample size under D2.

With these specifications, the total number of patients enrolled can be calculated as $(k-m)n_{i1} + mn_{i2}$. We further explore the design for values of $\alpha_t$ and $\alpha_{post}$ of $(0.05, 0.1, 0.15, 0.2, 0.25, 0.3, 0.35, 0.4)$, setting these two parameters independently. For each setting, we generate $10000$ simulated trials for the evaluation and comparison.

In order to assess the performance under various scenarios, we assess multiple outcome measurements (Supplemental Table S1). In addition to the measurements of family-wise error rate, power, and efficiency, other measurements include the coverage of the confidence interval for HR and estimation bias for HR. Specifically, the CI coverage for HR is defined as the probability that the estimated $95\%$ CI for the HR covers the true HR given that the individual test passed for that indication. Representing the relative difference in the true HR and the estimated HR, the estimation bias of HR is defined as the ratio of estimated average HR and true pooled HR for those indications that pass the individual tests minus $1$. To evaluate the outcome measurements, we calculated the average value for each measure over the $10000$ simulations.

### SUPPLEMENTAL TABLES

Table S1. Glossary of terms for simulation study

| Design parameters | Value(s) in simulation study | Descriptions |
|---|---|---|
| The state of nature | | |
| $g$ | $0, \dots, k$ | The number of active indications at the beginning |
| $\theta_i$ | At null value $\theta_0 = 1$,<br>At active value $\theta_1 = 0.5,0.6,0.7,0.8$. | The hazard ratio of experimental arm vs. control arm for the definitive endpoint of each indication |
| Study design input parameters | | |
| $k$ | $3,4,5,6$ | The number of tumor indications at the beginning |
| D | D1, D2, D3 | Sample size adjustment strategies |
| $t$ | $0.5$ | A common information time for the interim analysis |
| $\alpha_t$ | $0.05,0.1,0.15,0.2,0.25,0.3,0.35,0.4$ | A common bar for the interim analysis to prune inactive indications |
| $\alpha$ | $0.025$ | The false positive rate for pooled analysis together with the pruning step with respect to the global null hypothesis, after inflation from the nominal value $\alpha^*$ |
| $\beta$ | $0.025,0.05,0.1,0.2$ | False negative rate for the pooled analysis |
| $\alpha_{post}$ | $0.05,0.1,0.15,0.2,0.25,0.3,0.35,0.4$ | A common bar for the post - individual test, which is varied independently of $\alpha_t$ |
| $\alpha_{ref}$ | $0.025$ | False positive rate for the reference design |
| $\beta_{uncorrected}$ | $0.1$ | False negative rate for the uncorrected reference design |
| $S$ | $10000$ | The number of simulated replications |
| Calculated parameters | | |
| $n$ | $n = \frac{4(Z_{1-\alpha}+Z_{1-\beta})^2}{k(log\theta)^2}$ | Planned sample size for each tumor indication |
| $\alpha^*$ | Calculated by numerically solving the equation $\sum_{i=1}^{k} c(k,m)Q_0(\alpha^*,\alpha_t,m) = \alpha$ | The adjusted nominal level $\alpha^*$, at which the pooled analysis is nominally set to control the false positive rate $\alpha$ for the pooled |

| | | |
|---|---|---|
| | | analysis combined with the pruning step. |
| $\beta_{corrected}$ | $1-$power by indication corresponding to the given input parameters as determined by simulation | False negative rate for the corrected reference design |
| Simulated parameters | Descriptions | |
| $m$ | The number of tumor indications included in the pooled analysis | |
| $n_{i1}$ | Sample size for each tumor indication at the interim analysis ($nt$) | |
| $n_{i2}$ | Sample size for each tumor indication at the pooled analysis | |
| $Y_{i1}$ | The standardized test statistics for the $i$-th tumor indication at the interim analysis | |
| $Y_{i2}$ | The standardized test statistics for the $i$-th tumor indication at the final analysis | |
| $V_m$ | The standardized test statistics pooled from $Y_{i2}$ | |
| $d$ | The number of false positive indications passing the final individual tests | |
| $j$ | The number of true positive indications passing the final individual tests | |
| Outcome measurements | Estimates in simulation study | Descriptions |
| $\alpha_{net}$ (g) | $\frac{1}{S}\sum_{s=1}^{S} I\left(V_m^{(s)} > Z_{1-\alpha^{(s)}}\right) I\left(d^{(s)} > 0\right)$ | The probability of the basket trial passing the pooled test and at least one false positive indication passing the post-individual test for a given value of $g$. |
| Family-wise error rate (FWER) | $max\{\alpha_{net}(g): g = 0, \ldots, k\}$ | For any $g$ ($g = 0, \ldots, k$), the maximum probability of the basket trial passing the pooled test and at least one false positive indication passing the post-individual test |
| Power by indication | $\frac{1}{S}\sum_{s=1}^{S} \frac{j_{new}^{(s)}}{g} I\left(V_m^{(s)} > Z_{1-\alpha^{(s)}}\right)$ | The proportion of true positive indications that pass the post-individual test (requires that the pooled test passed) |
| Power by basket | $\frac{1}{S}\sum_{s=1}^{S}[I(g > 0) I\left(V_m^{(s)} > Z_{1-\alpha^{(s)}}\right)]$ | The probability of an active basket (one that contains at least one active indication) passing the pooled analysis |
| Sample size | $(k-m)n_{i1} + mn_{i2}$ | The sample size of a basket trial |
| Efficiency | $\frac{power \times g}{\frac{1}{S}\sum_{s=1}^{S}\left(\sum_{i=1}^{m^{(s)}} \mathrm{n}_{i2}^{(s)} + \sum_{i=m^{(s)}+1}^{k} \mathrm{n}_{i1}^{(s)}\right)} = \frac{\sum_{s=1}^{S} j_{new}^{(s)} I\left(V_m^{(s)} > Z_{1-\alpha^{(s)}}\right)}{\sum_{s=1}^{S}\left(\sum_{i=1}^{m^{(s)}} \mathrm{n}_{i2}^{(s)} + \sum_{i=m^{(s)}+1}^{k} \mathrm{n}_{i1}^{(s)}\right)}$ | The ratio of average number of active indications that passed the post-individual tests divided by the average sample size |
| Uncorrected reference efficiency | $\frac{g(1-\beta_{uncorrected})(log\theta_1)^2}{4k\left(Z_{1-\alpha_{ref}} - Z_{1-\beta_{uncorrected}}\right)^2}$ | The ratio of estimated number of active indications divided by the pre-defined total sample size in the reference study powered at 90%. |

| | | |
|---|---|---|
| Uncorrected relative efficiency | Efficiency/Uncorrected reference efficiency | The ratio of efficiency and uncorrected reference efficiency |
| Corrected reference efficiency | $\frac{g(1-\beta_{corrected})(log\theta_1)^2}{4k\left(Z_{1-\alpha_{ref}}-Z_{1-\beta_{corrected}}\right)^2}$ | The ratio of estimated number of true positive indications divided by the pre-defined total sample size in the reference study at the same power by indication observed in the simulation in the basket trial with corresponding parameters, investigating the same indications. |
| Corrected relative efficiency | Efficiency/Corrected reference efficiency | The ratio of efficiency and corrected reference efficiency |
| Coverage for hazard ratio (HR) | $\frac{1}{S}\sum_{s=1}^{S}[\sum_{i=1}^{m^{(s)}} I\left(Y_{i2}^{(s)} > Z_{1-\alpha_{post}}\right) I(\lvert Y_{i2}^{(s)} + \log(HR_i)\sqrt{\frac{n_{i2}^{(s)}}{4}}\rvert < Z_{0.975})]/[\sum_{i=1}^{m^{(s)}} I(Y_{i2}^{(s)} > Z_{1-\alpha_{post}})]$ | The probability that the estimated 95% CI for HR covers the true HR given the individual test passed |
| Bias of estimated HR | $\frac{1}{S}\sum_{s=1}^{S}\{\left[\frac{1}{m^{(s)}}\sum_{i=1}^{m^{(s)}} \exp\left(-Y_{i2}^{(s)}\sqrt{\frac{4}{n_{i2}^{(s)}}}\right) I\left(Y_{i2}^{(s)} > Z_{1-\alpha_{post}}\right)\right]/[\sum_{i=1}^{m^{(s)}} HR_i \frac{n_{i2}^{(s)} I\left(Y_{i2}^{(s)} > Z_{1-\alpha_{post}}\right)}{\sum_{i=1}^{m^{(s)}} n_{i2}^{(s)} I\left(Y_{i2}^{(s)} > Z_{1-\alpha_{post}}\right)}] - 1\}$ | The relative difference in the true HR and the estimated HR, defined as the ratio of estimated average HR and true pooled HR for those indications that pass the individual tests minus 1. |

Table S2. Simulation results. Each row summarizes the results of $10000$ simulations for a given scenario with input parameters: hazard ratio (HR), number of indications ($k$), $\beta$, $\alpha_t$ (α_t), $\alpha_{post}$ (α_post), number of active indications ($g$), and sample size adjustment strategies. The outcome measurements include: $\alpha_{net}$(α_net), power by indication, power by basket, mean sample size over simulations, efficiency, $95\%$ CI coverage for HR, bias, uncorrected reference efficiency, corrected reference efficiency, uncorrected relative efficiency, and corrected relative efficiency.

### SUPPLEMENTAL FIGURES

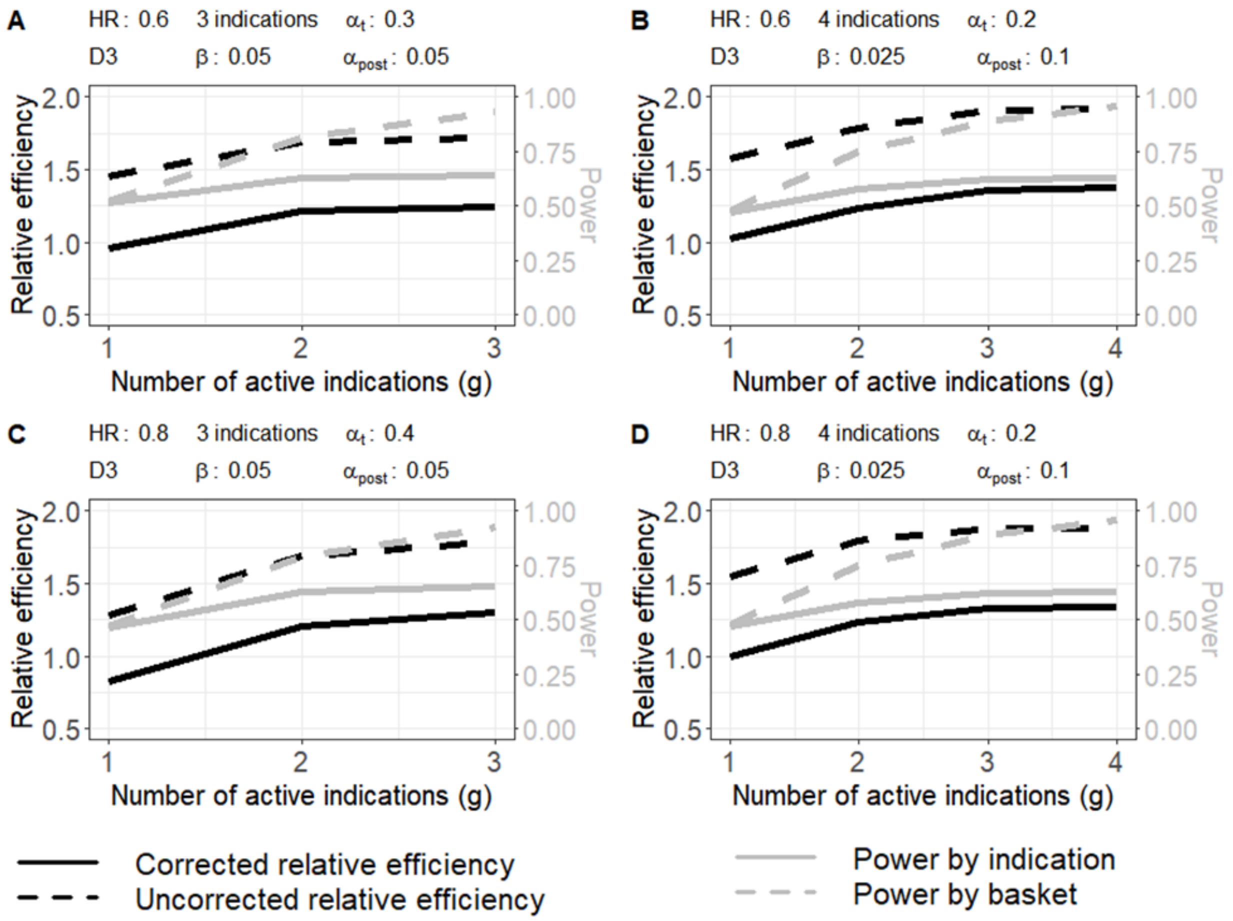


Figure S1. Recommended development approaches for (A) $3$ indications with $HR = 0.6$, (B) $4$ indications with $HR = 0.6$, (C) $3$ indications with $HR = 0.8$, and (D) $4$ indications with $HR = 0.8$. The X-axis represents the number of active indications (indications in which the drug provides clinical benefit), the primary Y-axis (left) represents the uncorrected/corrected relative efficiency, and the second Y-axis (right) represents the power by indication and by basket.

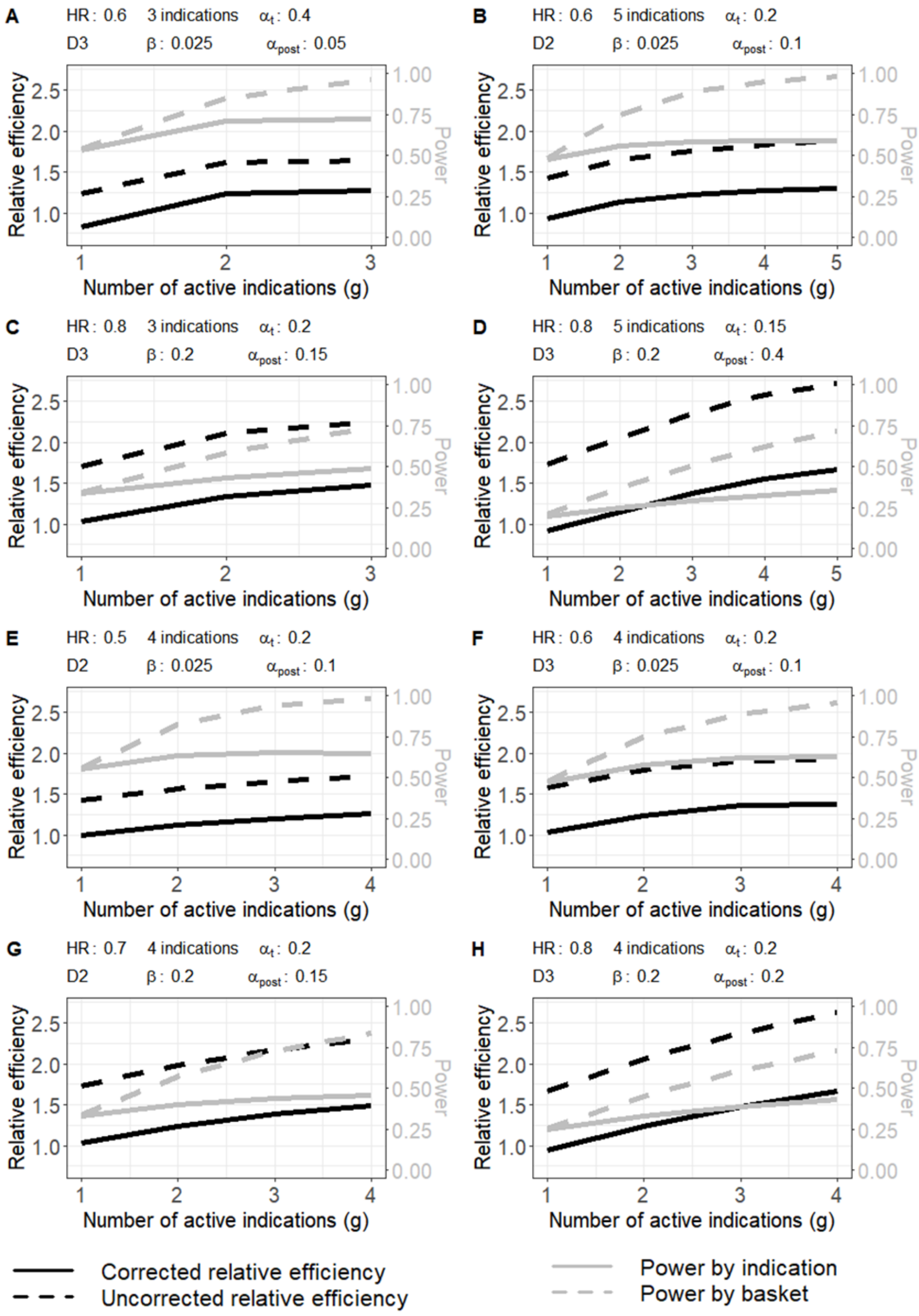
A
HR: 0.6 3 indications α_t: 0.4
D3 β: 0.025 α_post: 0.05
B
HR: 0.6 5 indications α_t: 0.2
D2 β: 0.025 α_post: 0.1
C
HR: 0.8 3 indications α_t: 0.2
D3 β: 0.2 α_post: 0.15
D
HR: 0.8 5 indications α_t: 0.15
D3 β: 0.2 α_post: 0.4
E
HR: 0.5 4 indications α_t: 0.2
D2 β: 0.025 α_post: 0.1
F
HR: 0.6 4 indications α_t: 0.2
D3 β: 0.025 α_post: 0.1
G
HR: 0.7 4 indications α_t: 0.2
D2 β: 0.2 α_post: 0.15
H
HR: 0.8 4 indications α_t: 0.2
D3 β: 0.2 α_post: 0.2
Relative efficiency
Power
Number of active indications (g)
Corrected relative efficiency
Uncorrected relative efficiency
Power by indication
Power by basket

Figure S2. Cases with maximum corrected relative efficiency for (A) $3$ indications with $HR = 0.6$, (B) $5$ indications with $HR = 0.6$, (C) $3$ indications with $HR = 0.8$, (D) $5$ indications with $HR = 0.8$, (E) $4$ indications with $HR = 0.5$, (F) $4$ indications with $HR = 0.6$, (G) $4$ indications with $HR = 0.7$, and (H) $4$ indications with $HR = 0.8$. The X-axis represents the number of active indications (indications in which the drug provides clinical benefit), the primary Y-axis (left) represents the uncorrected/corrected relative efficiency, and the second Y-axis (right) represents the power by indication and by basket.

### SUPPLEMENTAL R CODES

```
#Calculate alpha* given alpha_t, information time, and the number of indications.
#Reference: Cong Chen, Xiaoyun (Nicole) Li, Shuai Yuan, Zoran Antonijevic,
#Rasika Kalamegham & Robert A. Beckman(2016) Paper: Statistical Design and
#Considerations of a Phase 3 Basket Trial for Simutaneous Investigation of
#Multiple Tumor Types in  One Study, Statistics in  Biopharmaceutical Research,
#8:3, 248-257 DOI: 10.1080/19466315.2016.1193044. Function "mf2" is the original
#function from Chen's online code, it is used to express formula (3) in
#Chen-Beckman paper.

mf2 <- function(alphastar, alphat, t, m, k, d, Rho_for_endpoints=1) {
  # test1, test2, and test3 denotes correlation matrices for D1, D2, and D3.
  test <- matrix(0, ncol <- m + 1, nrow <- m + 1) # D1
  low <- rep(qnorm(1 - alphat), m + 1)
  low[m + 1] <- qnorm(1 - alphastar)
  up <- rep(Inf, m + 1)
  diag(test) <- 1

  for (i in 1:m)    test[i, m + 1] <- test[m + 1, i] <- switch (d,Rho_for_endpoints * sqrt(t / 
m),
                                                               Rho_for_endpoints * sqrt(t / k),
                                                               Rho_for_endpoints * sqrt(t / (k * (1 - t) + m * t)))

  # joint_probability1, joint_probability2, and joint_probability3 denotes the
  # joint probability in equation (3) in Chen-Beckman paper for D1, D2, and D3.
  joint_probability <- pmvnorm(lower <- low,upper <- up,mean <- rep(0, m + 1),corr <- 
test)[1]
```

```
  return(joint_probability)
}

# Function "type2" is the original function from Chen's online code, it is used
# to calculate alpha* by equation (3) in Chen-Beckman paper.
type2 <- function(alphastar, alphat, t, k, d, Rho_for_endpoints = .5) {
  # joint_probability denotes the joint probability in equation(3) in Chen-Beckman paper.
  joint_probability <- 0

  for (i in 1:k)     joint_probability <- joint_probability + factorial(k) / (factorial(i) * factorial(k
- i)) *
      (1-alphat)^(k-i)*mf2(alphastar=alphastar, alphat=alphat, t=t, m=i, k=k,
d=d,Rho_for_endpoints=Rho_for_endpoints)

  return(joint_probability - 0.025)
}

#"Simulation" function is denoted to calculate Type I error and power
# D denotes D1, D2, and D3, alpha_t denotes Type I error in interim stage,
# alpha_tt denotes Type I error after final stage for the post-trial test.
Simulation=function(alpha_t, alpha_tt, g, k, t = 0.5, design=c(D1=1,D2=2,D3=3),
                    Rho_for_endpoints,hr, delta=-log(hr), n, simulation_times=10000) {

  #set seed before simulation is to guarantee the results are reproductable.
  set.seed(123)
  # print(dummy_indication)
  #t denotes information time
  dummy_indication= sample(c(rep(1, g), rep(0, k-g)))
  delta=delta * dummy_indication; hr = exp(-delta)
```

```
  # print(delta);print(hr)
  #alphastar denotes alpha* under D1, D2 and D3 by Chen-Beckman's formula (2), the
  #function of "uniroot" is from Chen's online code.
  alphastar <- lapply(design,function(dd)
    uniroot(type2, c(0, 1), alphat=alpha_t, t=t, k=k,
d=dd,Rho_for_endpoints=Rho_for_endpoints)$root)

  mean_n <- delta * sqrt(n * t / 4)
  ###### Simulate result in before pruning
  x1 <- t(sapply(1:k,function(ln) if(dummy_indication[ln]==1)
{rnorm(simulation_times,mean=mean_n[ln], sd = 1)
    }else {rnorm(simulation_times, mean = 0, sd = 1)}))
  passed_pruning <- (x1 > qnorm(1 - alpha_t))

  #m denotes the number of remained indications after pruning
  m=colSums(passed_pruning)

  # Calculate sample size after pruning based on adjustment strategies.
  sample_size <- lapply(design,function(dd) apply(passed_pruning,2, function(pptmp)
    switch (dd,n * pptmp, ceiling(k * n / ifelse(sum(pptmp)>0,sum(pptmp),Inf)) * pptmp,
            ceiling((n * t + k * n * (1 - t) / ifelse(sum(pptmp)>0,sum(pptmp),Inf))) * pptmp)))

  ## calculate total sample size in the trial
  total_sample_size <- lapply(design,function(dd) apply(passed_pruning, 2, function(l)
    switch (dd,
            sum(n*l)+sum(n*t*(1-l)),
            ifelse(sum(l)>0,sum(n)+sum(n*t*(1-l)),sum(n*t*(1-l))),
            ifelse(sum(l)>0,sum(n),sum(n*t*(1-l)))
    )))
```

```
  # Calculate correlation between Yi1 and Yi2 if not all indications are pruned after
pruning step.
  rho_between_standardized_test_statistics <- lapply(design,function(dd) sapply(m,
function(mi)
    ifelse(mi>0, switch(dd,
                    Rho_for_endpoints * sqrt(t),
                    Rho_for_endpoints * sqrt(t * mi / k),
                    Rho_for_endpoints * sqrt(t / (t + k * (1 - t) / mi))
    ), 0)))

  # Generate Yi2 based on Yi1, corr(Yi1, Yi2), and adjusted sample size.
  #mean in the interim stage, mu1=sqrt(n*t/4)*delta*dummy_indication
  #representing means in the interim stage
  mu1 <- sqrt(n * t / 4) * delta * dummy_indication

  #mean in the final stage, mu2=sqrt(sample_size*1/4)*delta*dummy_indication
  #representing means in the final stage
  mu2 <- lapply(design, function(dd) (sqrt(sample_size[[dd]] / 4) * delta *
dummy_indication))

  #To generate Yi2, variance of Yi2 is sqrt((1-corr(Yi1,Yi2)^2))*s2^2,s2=1 in
  #our case(standardized normal distribution), given one of D1, D2, or D3.
  sd2 <- lapply(design, function(dd) sqrt((1 -
rho_between_standardized_test_statistics[[dd]] ^ 2)))

  #To generate Yi2, mean of Yi2(denoted as
  #mean_x2)=mu2+(s2/s1)*corr(Yi1,Yi2)*(Yi1-mu1),s1=s2=1 in our
  #case(standardized normal distribution) given one of D1, D2, or D3.
```

```
  mean_x2 <- lapply(design, function(dd)  mu2[[dd]] + (rep(1,k) %o%
rho_between_standardized_test_statistics[[dd]]) *
                      (x1 - mu1))

  #Generate Yi2 based on mean and variance for 3 indications, given one of D1,
  #D2, or D3.
  x2 <- lapply(design, function(dd) sapply(1:simulation_times, function(nsim)
    rnorm(mean_x2[[dd]][,nsim], mean_x2[[dd]][,nsim], sd = sd2[[dd]][nsim])))

  #Calculate V_m statistics, denotes as the sum of Yi2 for those indications
  #passed pruning at interim stage, divided by square root of number of
  #indications remained after pruning
  vm <- lapply(design, function(dd) sapply(1:simulation_times, function(nsim)
    ifelse(m[nsim]>0,sum(x2[[dd]][passed_pruning[,nsim],nsim]) / sqrt(m[nsim]),0)))

  # p_value_for_final_stage_testing denotes p-values for each simulation
  #time to compare with alphastar later
    p_value_for_final_stage_testing <- lapply(vm, pnorm)

  #If pooled indications are positive, do a post-check on individual
  #indications at alpha=post_alpha_t, and discard indications that do not
  #achieve statistical significance.
  #passed_pruning_post_trial denotes indications that passed pruning and pass
  #post-trial test
  passed_pruning_post_trial <-lapply(design, function(dd)
    (x1 > qnorm(1 - alpha_t) & x2[[dd]] > qnorm(1 - alpha_tt)))

  # calculate the coverage rate of 95% CI for individual indications and
  # pooled indications.
```

```
# coverage of individual approved indications
coveragebias=lapply(design, function(dd) sapply(1:simulation_times,function(nsim) {

  pppttmp=passed_pruning_post_trial[[dd]][,nsim];wttmp=n[pppttmp]/sum(n[pppttmp])
  pppttmp.tp=(passed_pruning_post_trial[[dd]][,nsim]) & (dummy_indication==1)
  wttmp.tp=n[pppttmp]/sum(n[pppttmp])

  hrtmp=exp(- x2[[dd]][,nsim] * sqrt(4 / sample_size[[dd]][,nsim]))[pppttmp]
  hrtmp.tp=exp(- x2[[dd]][,nsim] * sqrt(4 / sample_size[[dd]][,nsim]))[pppttmp.tp]

  c(individual=ifelse(sum(pppttmp)>0,
                    sum(abs(mu2[[dd]][pppttmp,nsim]-x2[[dd]][pppttmp,nsim])<
                          qnorm(p = 0.025, lower.tail = F))/sum(pppttmp),NA),

    pooled=ifelse(sum(pppttmp)>0,
                abs(sum(x2[[dd]][pppttmp,nsim]) / sqrt(sum(pppttmp))+
log(sum(hr[pppttmp]*wttmp))*sqrt(sum(sample_size[[dd]][pppttmp,nsim]) / 4))<
                  qnorm(p = 0.025, lower.tail = F),NA),

    bias1=ifelse(sum(pppttmp)>0,mean(hrtmp)/ mean(hr[pppttmp])-1,NA),
    bias2=ifelse(sum(pppttmp)>0,sum(hrtmp * wttmp) / sum(hr[pppttmp] * wttmp)-1,NA),

    indiv.tp=ifelse(sum(pppttmp.tp)>0,
                    sum(abs(mu2[[dd]][pppttmp.tp,nsim]-x2[[dd]][pppttmp.tp,nsim])<
                          qnorm(p = 0.025, lower.tail = F))/sum(pppttmp.tp),NA),
    pooled.tp=ifelse(sum(pppttmp.tp)>0,
                  abs(sum(x2[[dd]][pppttmp.tp,nsim]) / sqrt(sum(pppttmp.tp))+
```

```
log(sum(hr[pppttmp.tp]*wttmp.tp))*sqrt(sum(sample_size[[dd]][pppttmp.tp,nsim]) / 4))<
                    qnorm(p = 0.025, lower.tail = F),NA),
      bias1.tp=ifelse(sum(pppttmp.tp)>0, mean(hrtmp.tp)/ mean(hr[pppttmp.tp])-1,NA),
      bias2.tp=ifelse(sum(pppttmp.tp)>0,sum(hrtmp.tp * wttmp.tp) / sum(hr[pppttmp.tp] *
wttmp.tp)-1,NA)
    )}))

  #Record tp, fp in each simulation time after interim stage
  #tp denotes the number of active remained indication after pruning, fp denotes the
number of remained
  #indication after pruning.

  fp= lapply(design,function(dd) colSums(dummy_indication==0 &
passed_pruning_post_trial[[dd]]==1))

  #j, denotes the number of active indications remained after pruning and pass
  #post-trial test
  #(if we don't do post trial test, we change passed_pruning_post_trial to
  #passed_pruning in this line.)
  tp=lapply(design, function(dd)  colSums(dummy_indication==1 &
passed_pruning_post_trial[[dd]]==1))

  # Use the formula of Type I error and powers to get simulation results.

  final_pooled_test=lapply(design, function(dd)
    m > 0 & p_value_for_final_stage_testing[[dd]] > (1 - alphastar[[dd]]))

  fptp <-lapply(design, function(dd)
    c(type_I_error=sum(final_pooled_test[[dd]] & fp[[dd]] > 0) / simulation_times,
```

```
    power1=ifelse(g>0,sum(tp[[dd]][final_pooled_test[[dd]]]) / (g * simulation_times),0),
    power2=ifelse(g>0,sum(final_pooled_test[[dd]])/simulation_times,0)))

  ## Average total sample size
  average_total_sample_size <- lapply(total_sample_size, mean)

  ## Efficiency
  efficiency <- lapply(design, function(dd) c(efficiency=g * fptp[[dd]]['power1'] /
average_total_sample_size[[dd]]))
  inverse_efficiency <- lapply(design, function(dd) 1/efficiency[[dd]])

  ## Coverage and bias
  coveragebiasmean <-lapply(coveragebias,rowMeans,na.rm=TRUE)
  cbmse <-lapply(coveragebias,function(cb) apply(cb, 1, function(cbtmp)
    mean((cbtmp-mean(cbtmp,na.rm = TRUE))^2,na.rm = TRUE)))

  output <- list(test=fptp, mean_samplesize=average_total_sample_size,
efficiency=efficiency,cbmse=cbmse,
                 mean_coveragebias=coveragebiasmean,coveragebias=coveragebias)

  return(output)
}
```
